# Autoantibodies Against Ro/SS-A, CENP-B, and La/SS-B are Increased in Patients with Kidney Allograft Antibody-Mediated Rejection

**DOI:** 10.1101/2020.12.02.408922

**Authors:** Sergi Clotet-Freixas, Max Kotlyar, Caitriona M. McEvoy, Chiara Pastrello, Sonia Rodríguez-Ramírez, Sofia Farkona, Heloise Cardinal, Mélanie Dieudé, Marie-Josée Hébert, Yanhong Li, Olusegun Famure, Peixuen Chen, S. Joseph Kim, Emilie Chan, Igor Jurisica, Rohan John, Andrzej Chruscinski, Ana Konvalinka

## Abstract

Antibody-mediated rejection (AMR) causes >50% of late kidney graft losses. Although donor-specific antibodies (DSA) against HLA cause AMR, antibodies against non-HLA antigens are also linked to rejection. Identifying key non-HLA antibodies will improve our understanding of antibody-mediated injury.

We analyzed non-HLA antibodies using protein microarrays in sera from 91 kidney transplant patients with AMR, mixed rejection, acute cellular rejection (ACR), or acute tubular necrosis (ATN). IgM and IgG antibodies against 134 non-HLA antigens were measured pre-transplant, at the time of biopsy-proven diagnosis, and post-diagnosis. Findings were validated in 60 kidney transplant patients from an independent cohort.

Seventeen non-HLA antibodies were significantly increased (p<0.05) in AMR and mixed rejection compared to ACR or ATN pre-transplant, nine at diagnosis and six post-diagnosis. AMR and mixed cases showed significantly increased pre-transplant levels of IgG anti-Ro/SS-A and anti-CENP-B, compared to ACR. Together with IgM anti-CENP-B and anti-La/SS-B, these antibodies were also significantly increased in AMR/mixed rejection at diagnosis. Increased IgG anti-Ro/SS-A and anti-CENP-B pre-transplant and at diagnosis, and IgM anti-La/SS-B at diagnosis, were associated with the presence of microvascular lesions, but not with tubulitis or interstitial/total inflammation. All three antibodies were associated with the presence of class-II DSA (p<0.05). Significantly increased IgG anti-Ro/SS-A in AMR/mixed compared to ACR (p=0.01), and numerically increased IgM anti-CENP-B (p=0.05) and anti-La/SS-B (p=0.06), were validated in the independent cohort.

This is the first study that implicates autoantibodies against Ro/SS-A and CENP-B in AMR. These non-HLA antibodies may participate in the crosstalk between autoimmunity and alloimmunity in kidney AMR.

**SIGNIFICANCE STATEMENT:** Antibody-mediated rejection (AMR) causes >50% of kidney graft losses. Although donor-specific antibodies against HLA cause AMR, antibodies against non-HLA antigens are also linked to rejection. Serum samples of 91 kidney transplant patients were analyzed using protein arrays against 134 non-HLA antigens. AMR and mixed rejection cases showed significantly increased pre-transplant levels of IgG anti-Ro/SS-A and anti-CENP-B, compared to acute cellular rejection. Together with IgM anti-CENP-B and anti-La/SS-B, these antibodies were significantly increased in AMR/mixed rejection at diagnosis and were validated in a second, independent cohort. Increased IgG anti-Ro/SS-A, IgG anti-CENP-B and IgM anti-La/SS-B were associated with the presence of microvascular lesions and anti-HLA class-II antibodies. This is the first study to implicate anti-Ro/SS-A, anti-La/SS-B and anti-CENP-B autoantibodies in AMR.

## INTRODUCTION

Antibody-mediated rejection (AMR) causes >50% of late graft failures in kidney transplantation.^1^ AMR is usually caused by antibodies against human leukocyte antigens (HLA). Although histologic findings suggestive of AMR (i.e., microvascular inflammation) are linked to anti-HLA– mediated injury, some patients develop these lesions in the absence of anti-HLA donor-specific antibodies (DSA).^2–6^ In turn, not all transplant patients with anti-HLA DSA develop rejection,^7^ suggesting the involvement of other mechanisms in AMR.

Non-HLA allo- or autoantibodies may contribute to the pathogenesis of AMR. Antibodies against vimentin,^8,9^angiotensin-II type-1 receptor (AT1R),^2,10–15^collagen,^16,17^fibronectin,^16^perlecan/LG3,^18–21^and agrin,^5^ as well as anti-apoptotic cell autoantibodies,^22–25^ are associated with reduced survival and allograft rejection.^2,18,20,26^ Non-HLA antibodies are not routinely monitored and their contribution to kidney allograft injury is unclear. Moreover, their dynamic levels and their relationship with cellular rejection or other forms of graft injury remain unknown.

Production of autoantibodies may relate to viral infections, molecular mimicry, cryptic antigen exposure,^27–32^ or as yet unrecognized mechanisms. Autoantibodies produced post-transplant could result from immunotherapy-induced loss of regulatory T-cell proliferation and loss of tolerance to self-antigens.^33,34,35^ While several non-HLA autoantibodies recognized in systemic lupus erythematosus (SLE) and connective tissue disease have been extensively studied in autoimmunity,^36–39^ their role in alloimmunity has not been examined. Yet, autoimmune and alloimmune kidney injury share similarities, especially with regards to vascular injury.^40,41^ Furthermore, both SLE and allograft rejection^42^ are characterized by Th17 responses.^43,44^ There is increasing recognition of the interplay between allo- and autoimmunity,^17,23,45^ and this crosstalk may perpetuate injury.^46^

Our aim was to identify non-HLA antibodies associated with AMR, and to determine their evolution over time and their link to DSA and histopathology lesions. We describe herein a retrospective cohort of 91 kidney transplant patients with 134 non-HLA antibodies measured pre-transplant, at the time of indication biopsy-based diagnosis, and post-diagnosis. Antibodies previously implicated in solid organ transplant injury or autoimmunity were measured using protein arrays. We identified anti-Ro/SS-A(52KDa), anti-CENP-B, and anti-La/SS-B antibodies as significantly increased in kidney transplant recipients with AMR, compared to patients with acute cellular rejection (ACR) or acute tubular necrosis (ATN). These antibodies were associated with class-II DSA and microvascular lesions. We validated these antibodies in an external, independent cohort. This is the first study, to our knowledge, that links autoimmunity-related antibodies to humoral alloimmunity.

## METHODS

### Study Design and Patient Population

We studied two groups of patients, one for discovery and one for validation. For the discovery phase, we identified kidney transplant recipients at the University Health Network (UHN) in Toronto, with rejection diagnosed between 2008 and 2016, by searching the CoReTRIS registry.^47^ We selected cases with a histological diagnosis of rejection on a for-cause biopsy, and at least one serum sample available in the HLA laboratory. Samples with insufficient volume and/or retrieved within 21 days after the patient received plasmapheresis (PLEX) and/or intravenous immunoglobulin infusion (IVIG) were excluded from further study. Patient exclusion criteria were 1) no serum sample available post-transplant; and 2) all serum samples affected by PLEX and/or IVIG. Finally, we selected cases with ATN that were graft-age matched to AMR and to ACR cases. A renal pathologist (R.J.) scored the biopsies according to the Banff classification (2017).^3^ Twenty-eight of these 91 patients were described in our recent study.^48^ This study was approved by the UHN institutional research ethics board (CAPCR identifier 13-7261).

Serum samples were collected pre-transplant, ‘at diagnosis’ (within 30 days of the indication biopsy date), and ‘post-diagnosis’ (collected >30 days after the indication biopsy). The time intervals between the indication biopsy and the post-diagnosis serum collection ranged from 37 days to 2,193 days. The presence of anti-HLA class-I and class-II antibodies in the sera was assessed using Luminex single-antigen bead assays, as part of standard clinical practice. To assess non-HLA antibody levels, we quantified IgG and IgM antibodies against 134 non-HLA antigens, using a VersArray Chipwriter Pro antigen microarray platform (Virtek, Canada). Antigen characteristics are described in **Table S1**.

For external validation, sera from 60 kidney transplant patients from Center Hospitalier de l’Université de Montréal (CHUM, Montreal) were retrieved ‘at diagnosis’ (median time of ≤4 days from the indication biopsy). In these samples, IgG and IgM levels against the key non-HLA antibodies identified in the discovery study, were measured using the same microarray platform. The validation cohort was part of a previous study of LG3-related antibodies, and consisted of 29 stable non-rejecting cases, 16 cases with ACR, and 15 cases with acute vascular rejection (AVR), as described previously.^49^ Upon review of the AVR cases, we were able to assign a diagnosis of AMR or mixed rejection (n=5), ACR grade 2-3 (n=3), and AVR with insufficient information to delineate between AMR and ACR (n=7). The study was approved by the clinical research ethics committee at CHUM (Research Ethics Board number 2008-2545, HD.07.034).

### Histopathology

Each indication biopsy was embedded in paraffin and 3μm sections were obtained in a microtome (Leica). Sections were then deparaffinized through graded alcohols and subjected to hematoxylin/eosin, trichrome, periodic acid–Schiff, and periodic Schiff-methenamine stains; and examined under a light microscope. Staining of C4d was also performed on additional 4μm frozen sections by immunofluorescence. Morphologic features were diagnosed and given a semiquantitative score (0-3) by R.J., according to the updated Banff classification.^3,48^

### Non-HLA Antigen Microarrays

#### Antigen Library and Microarray Generation

The 134 antigens including proteins, peptides, and cell lysates were diluted to 0.2 mg/ml in PBS and stored in aliquots at −80°C. These antigens were selected because of their importance in autoimmune diseases,^50,51^ or because they were linked to humoral rejection of several organs including kidney,^52^ lung^53^ or heart.^54^ To generate the protein microarrays, the 134 antigens screened in this study were spotted in duplicate onto two-pad FAST nitrocellulose coated slides (Maine Manufacturing, Sanford, ME) using a VersArray Chipwriter Pro microarrayer (Virtek, Toronto, Canada) as previously described.^54–56^ Slides were arrayed using solid pins (Arrayit, Sunnyvale, CA), which generated antigen spots with a diameter of approximately 500μm. This process was conducted at 55% relative humidity and room temperature. Two microarrays were generated on each slide. Nine additional empty spots were measured as blank, to determine the background fluorescence. In both sets of microarrays (discovery and validation), PBS was spotted as negative control, while human IgG (whole molecule and Fc fragment) and human IgM (whole molecule) were spotted to confirm specificity of the anti-IgG and anti-IgM secondary antibodies, respectively.

#### Sample processing

The antigen microarray platform was used to screen for IgG and IgM against the 134 non-HLA antigens as previously described.^54^ Slides were first dried at room temperature and placed in FAST frames (Maine Manufacturing). Each frame supported a total of 4 slides (8 protein arrays). Microarrays were then incubated overnight with 700μL of blocking buffer (PBS, 5% FBS, 0.1% Tween) at 4°C. The following day, blocking buffer was removed and each microarray was incubated for 1 hour at 4°C with 600μL of a different serum sample at a dilution of 1:100 in blocking buffer. A serum sample from a patient with SLE with known pronounced IgG reactivity against Ribo P0, Ribo P1 and Ribo P2, as well as high IgM reactivity against ssDNA and dsDNA, was arrayed as a quality control. Blocking buffer alone was used as a negative control. Microarrays were then washed 3 times for 10 minutes with PBST (PBS and 0.1% Tween) at room temperature, and subsequently probed with secondary antibodies for 45 minutes at 4°C. Slides were probed with a mixture of secondary antibodies consisting of Cy3-labeled goat anti-human IgG (Jackson ImmunoResearch, West Grove, PA) at a dilution of 1:2,000 and Cy5-labeled goat anti-human IgM (Jackson ImmunoResearch) at a dilution of 1:1,000. Microarrays were washed again 3 times for 10 minutes with PBST, and slides were dried by centrifugation at 220G for 5 minutes at room temperature. Slides were kept at room temperature and protected from the light until scanning.

To minimize the potential effect of frame-to-frame or day-to-day variability when comparing different study groups, all groups were proportionally represented in each processing day. Samples from the same patient collected at different time points were analyzed in the same frame, to avoid potential batch effects when generating time-course data. Toronto (discovery) and Montreal (validation) cohorts could not be directly compared, since samples were analyzed on different days, and arrays are subject to batch effects.

#### Quantification of fluorescence intensity

The fluorescent signals of Cy3 (ʎ=532 nm) and Cy5 (ʎ=635 nm) were measured on an Axon 4200A microarray scanner (Molecular Devices, Sunnyvale, CA) using a Genepix 6.1 software (Molecular Devices). For each spot, the software calculated the median fluorescent intensity (MFI) minus the local background on the Cy3 and Cy5 channels. To confirm adequate background correction, nine blank spots were also measured in each array. The MFI of each antigen was then determined by averaging the intensity of the duplicate spots. Values of MFI ≥200 in at least one sample were set as detection threshold to further study the antibody signal against a particular antigen. A signal with MFI ≥200 was >2 SD from the average MFI of the blanks. As expected, the highest IgG signal across all samples was found at the antigen spots with human IgG (average MFI = 58,212.59) or human IgG Fc (average MFI = 64,363.63), while the highest IgM signal belonged to the spots with human IgM (average MFI = 51,100.25).

### Data Analysis and Bioinformatics

#### Assessment of data distribution

We evaluated the distributions of all IgG and IgM MFI values in the different diagnoses, and between the sexes. In each patient, a value of zero was given to all antibodies with MFI <200. The log2-transformed MFI values of each antibody in each study group were then used to create density plots using ggplot2 3.3.2^57^ in R 4.0.2.^58^ Log2-transformed values have been used in subsequent analyses.

#### Differential antibody levels between types of rejection

Differential antibody levels analysis of both data sets (discovery and validation) was performed in R. The Wilcoxon–Mann–Whitney nonparametric test was used to assess differences in non-HLA antibody levels between groups. By definition, both AMR and mixed cases show DSA and/or histological signs of antibody-mediated injury. In addition, the distributions of the IgG and IgM intensity values were comparable between these two forms of rejection. We thus combined the AMR and the mixed cases in one single ‘AMR/mixed’ group in the differential antibody MFI analysis, and compared this group to ACR and ATN. This enabled us to enhance the statistical power of the comparative analysis. Data are presented as medians ± SEM. P<0.05 was considered significant.

#### Antibody changes over time and hierarchical clustering analysis

Log2-transformed MFI values of the non-HLA antibodies with significantly different levels in AMR/mixed patients compared to ACR or ATN were used to generate violin plots of MFI distribution, to visualize antibody changes among groups at different time points. Scatter plots were used to compare MFI trends between the “pre-transplant” and “at diagnosis” time points. Plots were created using ggplot2. Values were then normalized using the formula (x-min(x))/(max(x)-min(x)) and used to build heatmaps with pheatmap 1.0.12^59^. Unsupervised hierarchical clustering of antibodies and samples was performed using pheatmap with default settings.

#### Association between non-HLA antibody levels and clinical variables

Associations between the MFI of non-HLA antibodies (continuous variable) and ordinal histopathology variables based on a semiquantitative 0-3 score (namely vascular fibrous intimal thickening, tubulitis, total inflammation, peritubular capillaritis, intimal arteritis, interstitial inflammation, glomerulitis, chronic glomerulopathy, and C4d deposition), were analyzed. Each histopathology variable was split into low (0 or 0-1) and high (1-3 or 2-3) categories, and Wilcoxon–Mann–Whitney nonparametric tests were used to assess differences in non-HLA antibody expression between these two categories. Associations between the MFI of non-HLA antibodies and categorical variables (namely presence=1 versus absence=0 of anti-HLA DSA, anti-HLA class-I DSA and anti-HLA class-II DSA), were also assessed by Wilcoxon–Mann– Whitney nonparametric tests. All analyses were conducted in R. P<0.05 was considered significant. The most significant associations between all clinical variables and non-HLA antibodies were represented using bubble plots (R and ggplot2).

#### Protein-protein interaction and network analysis

Physical protein-protein interactions of key non-HLA antibody targets were collected using the Integrated Interactions Database (version 05-2020, http://iid20.ophid.utoronto.ca, https://doi.org/10.1093/nar/gky1037)^60^. Interactions experimentally validated or predicted were retained and visualized using NAViGaTOR 3.0.13 (http://ophid.utoronto.ca/navigator). The gene names of the antibody targets that we previously found to be differentially expressed at the protein level in the AMR glomeruli or tubulointerstitium^48^ are also shown in the network.

## RESULTS

### Study Population

We studied sera from 43 patients with AMR, 20 patients with mixed rejection, 16 patients with ACR, and 12 patients with ATN (**Table 1**, **Fig. 1A**). Most patients were males, except for AMR cases, who had ~50% males. Patients with ATN were significantly older than other groups. The median time between transplantation and diagnostic biopsy was similar in patients with AMR, ACR, and ATN (9.5-15.5 days), but higher among mixed cases (174 days). Most patients with AMR (86%) or mixed rejection (80%) had class-I and/or class-II DSA (**Table 1**). None of the ACR cases had DSA. Although 4/12 ATN patients had DSA, their biopsies did not show signs of rejection (**Table 2**). Glomerulitis and C4d deposition were detected exclusively among AMR and mixed cases. These two groups showed the highest scores for peritubular capillaritis. The highest interstitial inflammation, tubulitis, and total inflammation were observed in mixed rejection and ACR. Chronic glomerulopathy was found in one mixed and two AMR cases (**Table 2**).

**Table 1.** Clinical parameters of the patient cohort. *Autoimmune conditions included: primary sclerosing cholangitis (n=1), ANCA-vasculitis (n=4), SLE (n=2), and diabetes mellitus type I (n=2). ESKD, end stage kidney disease; FSGS, focal and segmental glomerulosclerosis; IQR, interquartile range; PCKD, polycystic kidney disease; SLE, systemic lupus erythematosus; TMA, thrombotic microangiopathy

|  | Antibody mediated rejection | Mixed rejection | Acute cellular rejection | Acute tubular necrosis |
| --- | --- | --- | --- | --- |
| **Number of patients** | 43 | 20 | 16 | 12 |
| **Sex - number of males, n (%)** | 22 (51) | 13 (65) | 12 (75) | 9 (75) |
| **Patient age at biopsy, median years (IQR)** | 50 (39, 58) | 48 (36.7, 56) | 44 (36.2, 51.5) | 64.5 (62.2, 69.7) |
| **Time post-transplant, median days (IQR)** | 12 (8, 22) | 174 (72, 737) | 15.5 (12, 29.2) | 9.5 (7.7, 12.2) |
| **Cause of ESKD** |  |  |  |  |
| Diabetic Nephropathy, n (%) | 9 (21) | 3 (15) | 1 (6.2) | 4 (33.3) |
| IgA Nephropathy, n (%) | 2 (4.6) | 2 (10) | 5 (31.2) | 1 (8.3) |
| PCKD, n (%) | 4 (9.3) | 5 (25) | 3 (18.7) | 2 (16.6) |
| Vasculitis, n (%) | 3 (7) | 0 (0) | 1 (6.2) | 0 (0) |
| FSGS, n (%) | 4 (9.3) | 2 (10) | 1 (6.2) | 1 (8.3) |
| Reflux, n (%) | 3 (7) | 0 (0) | 0 (0) | 0 (0) |
| Hypertension, n (%) | 3 (7) | 1 (5) | 0 (0) | 1 (8.3) |
| SLE, n (%) | 2 (4.6) | 1 (5) | 0 (0) | 0 (0) |
| TMA, n (%) | 2 (4.6) | 0 (0) | 0 (0) | 0 (0) |
| Unknown, n (%) | 3 (7) | 4 (20) | 0 (0) | 2 (16.6) |
| Other, n (%) | 8 (18.6) | 3 (15) | 5 (31.2) | 1 (8.3) |
| **Pre-existing autoimmune conditions\*, n (%)** | 7 (16.3) | 1 (5) | 1 (6.2) | 0 (0) |
| **Renal replacement therapy** | 40 (93) | 16 (80) | 14 (87.5) | 12 (100) |
| Intermittent hemodialysis, n (%) | 35 (81.3) | 11 (55) | 10 (62.5) | 11 (91.6) |
| Peritoneal dialysis, n (%) | 5 (11.6) | 5 (25) | 4 (25) | 1 (8.3) |
| Preemptive, n (%) | 3 (7) | 4 (20) | 2 (12.5) | 0 (0) |
| **Prior desensitization, n (%)** | 21 (48.8) | 3 (15) | 0 (0) | 0 (0) |
| Eculizumab, n (%) | 1 (2.3) | 0 (0) | 0 (0) | 0 (0) |
| Rituximab, n (%) | 3 (7) | 2 (10) | 0 (0) | 0 (0) |
| **Kidney transplant, donor type** |  |  |  |  |
| Living donor, n (%) | 21 (49) | 9 (45) | 9 (56.2) | 3 (25) |
| Deceased donor, n (%) | 22 (51) | 11 (55) | 7 (43.7) | 9 (75) |
| **Induction therapy** |  |  |  |  |
| Thymoglobulin, n (%) | 38 (88.3) | 16 (80) | 11 (68.8) | 12 (100) |
| Basiliximab, n (%) | 3 (7) | 3 (15) | 5 (31.2) | 0 (0) |
| Unknown, n (%) | 2 (4.6) | 1 (5) | 0 (0) | 0 (0) |
| **Maintenance therapy, number (%)** |  |  |  |  |
| Prednisone | 43 (100) | 20 (100) | 16 (100) | 12 (100) |
| Anti-proliferative | 42 (97.6) | 20 (100) | 16 (100) | 12 (100) |
| Calcineurin Inhibitor | 43 (100) | 20 (100) | 16 (100) | 12 (100) |
| **ABO incompatible, n (%)** | 5 (11.6) | 0 (0) | 0 (0) | 0 (0) |
| **Delayed graft function, n (%)** | 17 (39.5) | 3 (15) | 4 (25) | 2 (16.6) |
| **Primary non-function, n (%)** | 2 (4.6) | 1 (5) | 0 (0) | 0 (0) |
| **DSA, current or historic** |  |  |  |  |
| Any DSA | 37 (86) | 16 (80) | 0 (0) | 4 (33.3) |
| Class I, n (%) | 23 (53.4) | 7 (35) | 0 (0) | 4 (33.3) |
| Class II, n (%) | 31 (72) | 16 (80) | 0 (0) | 0 (0) |
| Unknown, n (%) | 2 (4.6) | 1 (5) | 1 (6.2) | 1 (8.3) |

**Figure 1.**
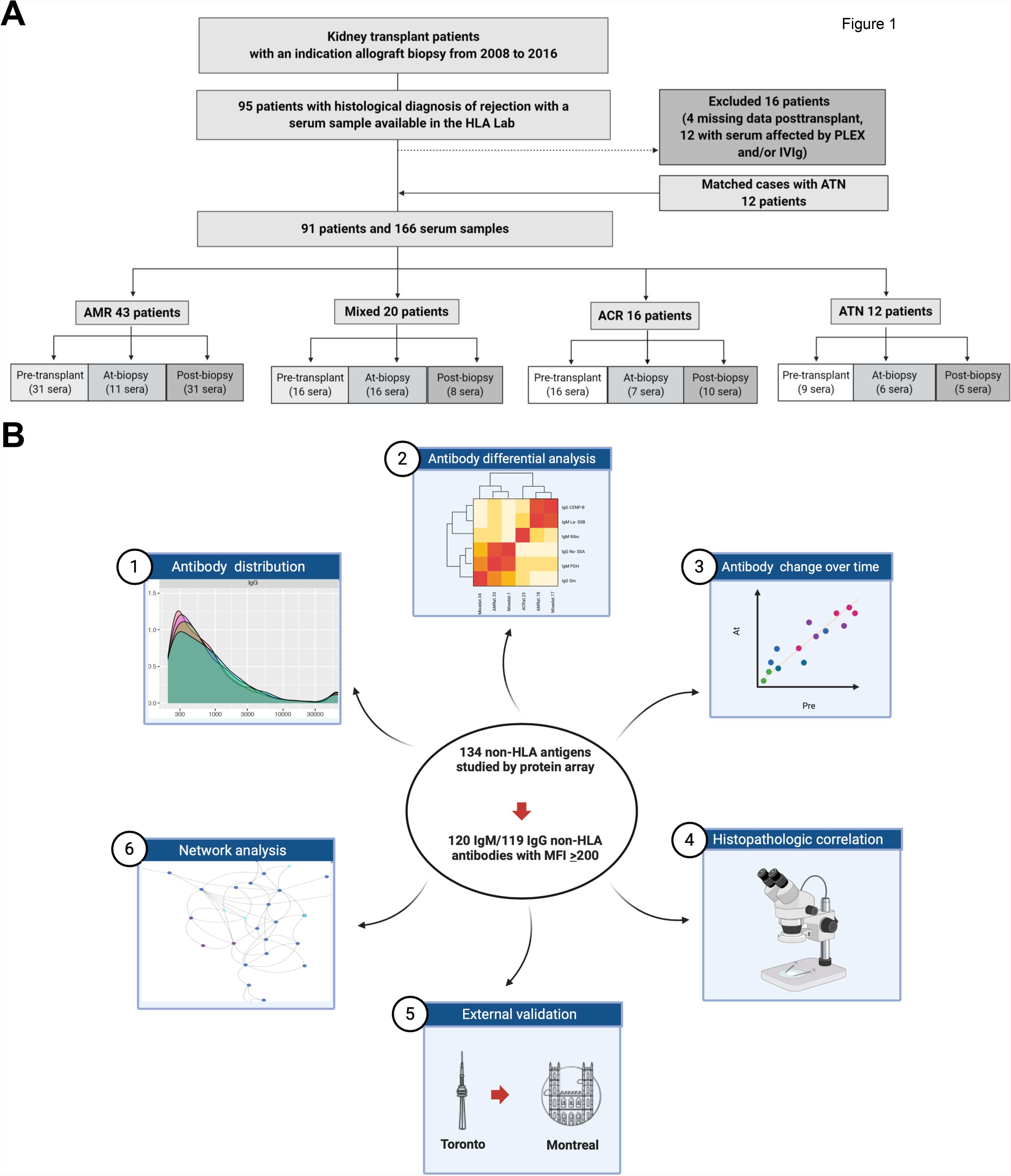
Experimental design and study workflow. In the discovery phase, we identified kidney transplant recipients with rejection diagnosed between 2008 and 2016 (A). Patient exclusion criteria were: 1) no serum sample available post-transplant; or 2) all serum samples collected within 21 days after PLEX and/or IVIG administration. Graft-age matched cases with acute tubular necrosis (ATN) were also included. A total of 166 sera were selected from 91 kidney transplant patients, with antibody-mediated rejection (AMR, n=43), ‘mixed’ antibody-mediated and cellular rejection (n=20), acute cellular rejection (ACR, n=16), or ATN (n=12). Our study workflow is shown in panel B. We first evaluated the MFI values of all the detected IgG and IgM, by comparing their distributions across different diagnoses. We then subjected all non-HLA antibodies to statistical analyses to assess differences between groups. We also performed clustering analyses to assess how the antibodies clustered in relation to the diagnoses and the anti-HLA DSA. Changes over time of key non-HLA antibodies were studied by plotting the MFI at diagnosis compared to MFI pre-transplant, in patients who had both samples available for the analysis. We next studied the association between the levels of each non-HLA antibody and the presence of histopathology lesions and/or anti-HLA DSA. We also interrogated our top antibodies of interest in an independent validation cohort. Finally, we built a protein-protein interaction network that integrates our top non-HLA antibody targets with our previous proteomics data sets of the AMR glomeruli and tubulointerstitium. DSA, donor-specific antibodies; Ig, immunoglobulin; MFI, median fluorescence intensity; PLEX, plasmapheresis; IVIG, intravenous immunoglobulin.

**Table 2.** Biopsy findings of the patient cohort. Histopathology lesions were evaluated according to the most updated Banff classification. IQR, interquartile range; ah, arteriolar hyalinosis; c4d, c4 deposition; cg, chronic glomerulopathy; ci, interstitial fibrosis; ct, tubular atrophy; cv, vascular fibrous intimal thickening; g, glomerulitis; i, interstitial inflammation; mm, mesangial matrix expansion; ptc, peritubular capillaritis; ti, total inflammation; t, tubulitis; v, intimal arteritis.

| **Median (IQR)** | **AMR (43)** | **MIXED (20)** | **ACR (16)** | **ATN (12)** |
| --- | --- | --- | --- | --- |
| **i** | 0 (0, 0) | 2 (1.7, 3) | 2 (1.7,2) | 0 (0, 0) |
| **t** | 0 (0, 0) | 2 (2, 3) | 3 (2.7, 3) | 0 (0, 0) |
| **ti** | 0 (0, 1) | 2 (2, 3) | 2 (2, 3) | 0 (0, 0) |
| **g** | 2 (0, 2) | 0 (0, 2) | 0 (0, 0) | 0 (0, 0) |
| **ptc** | 1 (0, 2) | 2 (1, 2) | 1 (0, 1) | 0 (0, 0) |
| **cg** | 0 (0, 0) | 0 (0, 0) | 0 (0, 0) | 0 (0, 0) |
| **mm** | 0 (0, 0) | 0 (0, 0) | 0 (0, 0) | 0 (0, 0) |
| **v** | 0 (0, 0) | 1 (0, 1) | 0 (0, 0) | 0 (0, 0) |
| **ci** | 0 (0, 0) | 1 (0, 1) | 1(0, 1) | 0 (0, 0) |
| **ct** | 0 (0, 0) | 1 (0, 1) | 1 (0.7, 1) | 0 (0, 1) |
| **ah** | 0 (0, 1) | 1 (0, 1) | 0 (0, 1) | 0 (0, 0) |
| **cv** | 0 (0, 1) | 1 (0, 1.5) | 1 (0, 1.7) | 1 (0, 1) |
| **C4d** | 3 (3, 3) | 3 (2, 3) | 0 (0, 0) | 0 (0, 0) |
| **Globally sclerosed glomeruli, % (IQR)** | 4 (0, 7) | 5 (0, 11.7) | 4 (0, 11.5) | 4 (0, 8.2) |

### The Distribution of IgG Antibodies is Conserved Across Time and Diagnoses

The workflow is shown in **Fig. 1B**. We analysed 166 serum samples using protein arrays against 134 non-HLA antigens (**Table S1**). We focused on 119 IgG and 120 IgM antibodies against non-HLA antigens detected with MFI≥200 in ≥1 sample. We examined the distributions of antibody levels in each group. The distribution of IgG antibodies was remarkably similar among groups and stable across time (**Fig. 2A-C**). Conversely, IgM antibody levels displayed a shift toward higher levels only in AMR and mixed rejection, at diagnosis (**Fig. 2B**). Male sex was associated with increased pre-transplant levels of IgG and IgM antibodies (**Fig. S1**).

**Figure 2.**
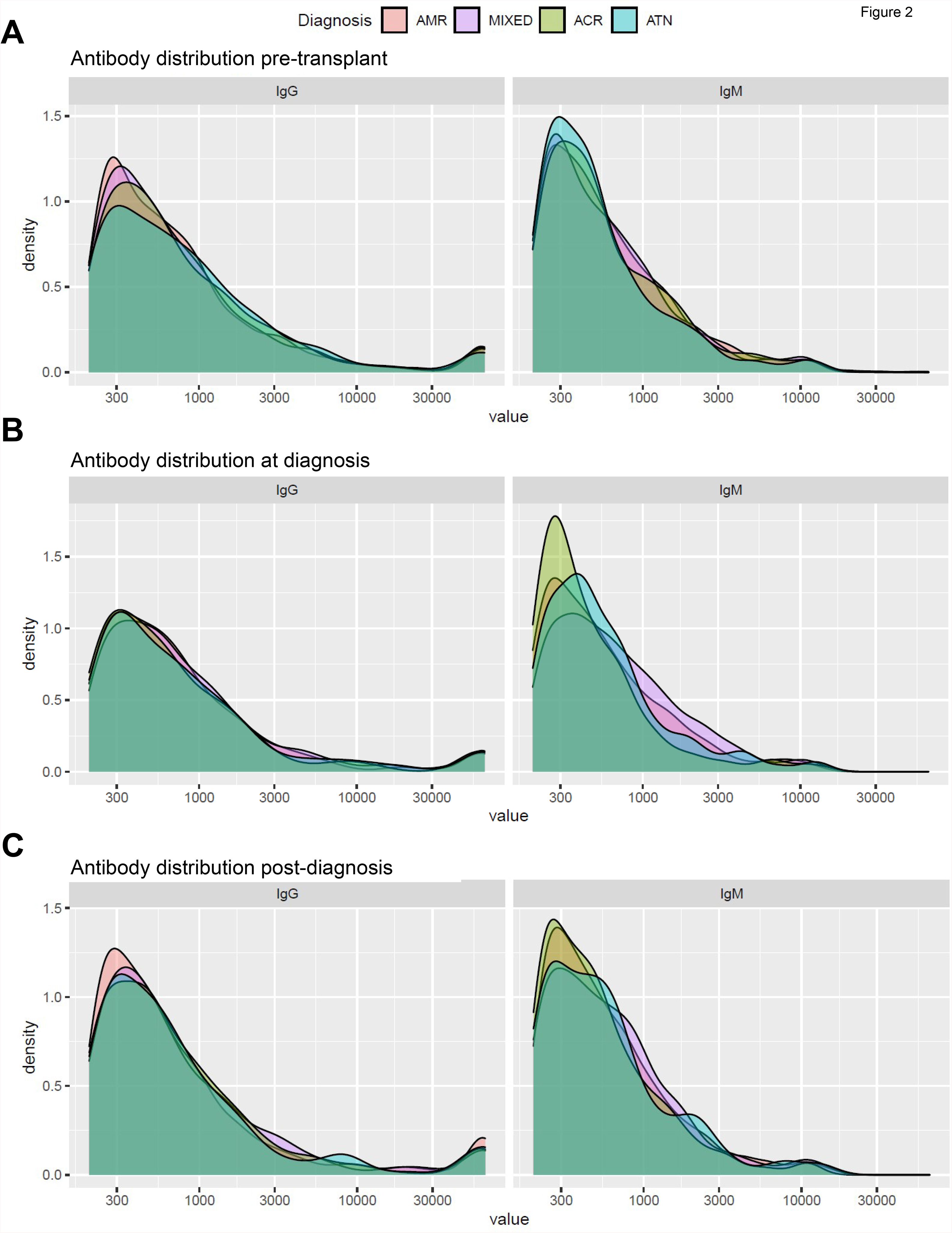
Distribution of antibody levels according to diagnosis. The density plots depict the distributions of IgG and IgM MFI values in AMR (orange), mixed rejection (purple), ACR (green), and ATN (teal) before transplant (A), at diagnosis (B), and post-diagnosis (C). In each patient, a value of zero was given to all antibodies with MFI < 200. Log2-transformed MFI values were used to create the plots using ggplot2 3.3.2 in R. The x-axis encompasses the range of all MFI values among the detected antibodies, while the y-axis represents the frequency (density) of each of these values. AMR, antibody-mediated rejection; ACR, acute cellular rejection; ATN, acute tubular necrosis; Ig, immunoglobulin; MFI, median fluorescence intensity.

### Antibodies Against Ro/SS-A, CENP-B, and La/SS-B are Increased in Kidney AMR and Mixed Rejection

We were predominantly interested in non-HLA antibodies associated with antibody-mediated injury. Interestingly, AMR and mixed rejection groups had analogous IgG and IgM antibody distributions, emphasizing their similarity (**Fig. 2**). Among the studied non-HLA antibodies, 36 were significantly altered (P<0.05) in AMR/mixed rejection compared to ACR and/or ATN, at one or more time points (**Table 3**), being 80% of them were increased in AMR/mixed rejection. IgG anti-Ro/SS-A(52KDa) and IgG anti-CENP-B were significantly increased in AMR/mixed patients compared to ACR, both pre-transplant and at diagnosis. Together with these 2 antibodies, IgM anti-CENP-B, IgM anti-La/SS-B, IgM anti-Ribo P1, and IgM anti-PDH were significantly increased at diagnosis in AMR/mixed patients, compared to ACR. Furthermore, the distributions of IgG anti-Ro/SS-A(52KDa), IgG and IgM anti-CENP-B, and IgM anti-La/SS-B, were remarkably similar between patients with AMR and mixed rejection, but different from ACR and ATN, and remained consistent across time (**Fig. 3A**). Hierarchical clustering showed that levels of IgG anti-Ro/SS-A(52KDa) pre-transplant and at diagnosis clustered with IgG and IgM anti-CENP-B, respectively, and were highest in patients with class-II DSA (**Fig.S2A,C**). Anti-mitochondrial antibodies against components of the pyruvate dehydrogenase complex, namely IgM anti-PDH and IgG anti-M2 (PCD-E2, OGDC-E2, BCOADC-E2 antigens), were also increased in AMR/mixed compared to ACR and ATN, at diagnosis (**Fig. 3A-B**, **Table 3**).

**Table 3.**
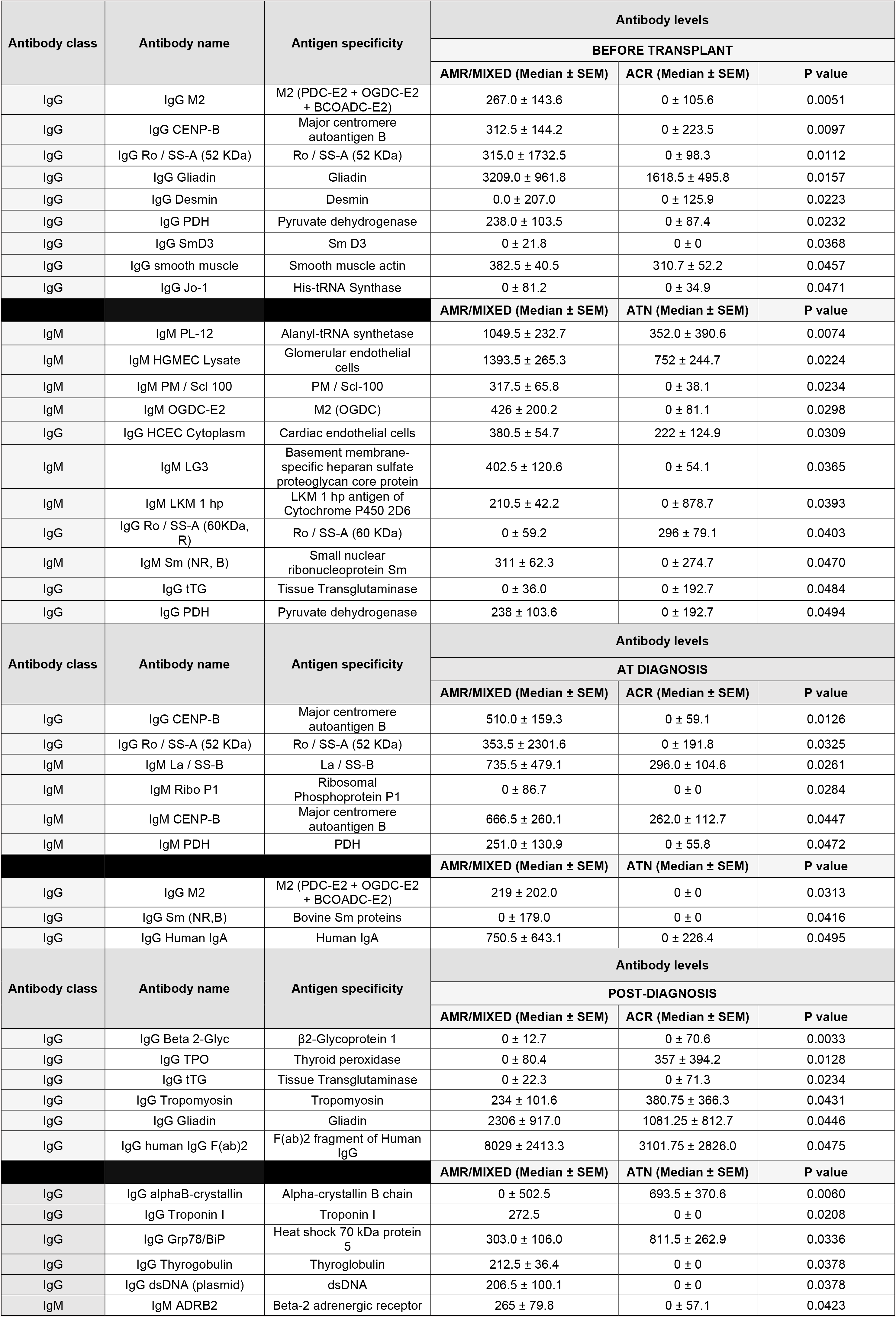
Antibodies against non-HLA antigens significantly altered in AMR/Mixed rejection, compared to ACR and/or ATN. Significantly altered (P value < 0.05) IgG and IgM antibodies before transplant, at diagnosis, and post-diagnosis are shown. Ig, immunoglobulin; AMR, antibody-mediated rejection; ACR, acute cellular rejection; ATN, acute tubular necrosis; SEM, standard error of the median; B, bovine; R, recombinant; NR, non-recombinant; hp, high purity.

**Figure 3.**
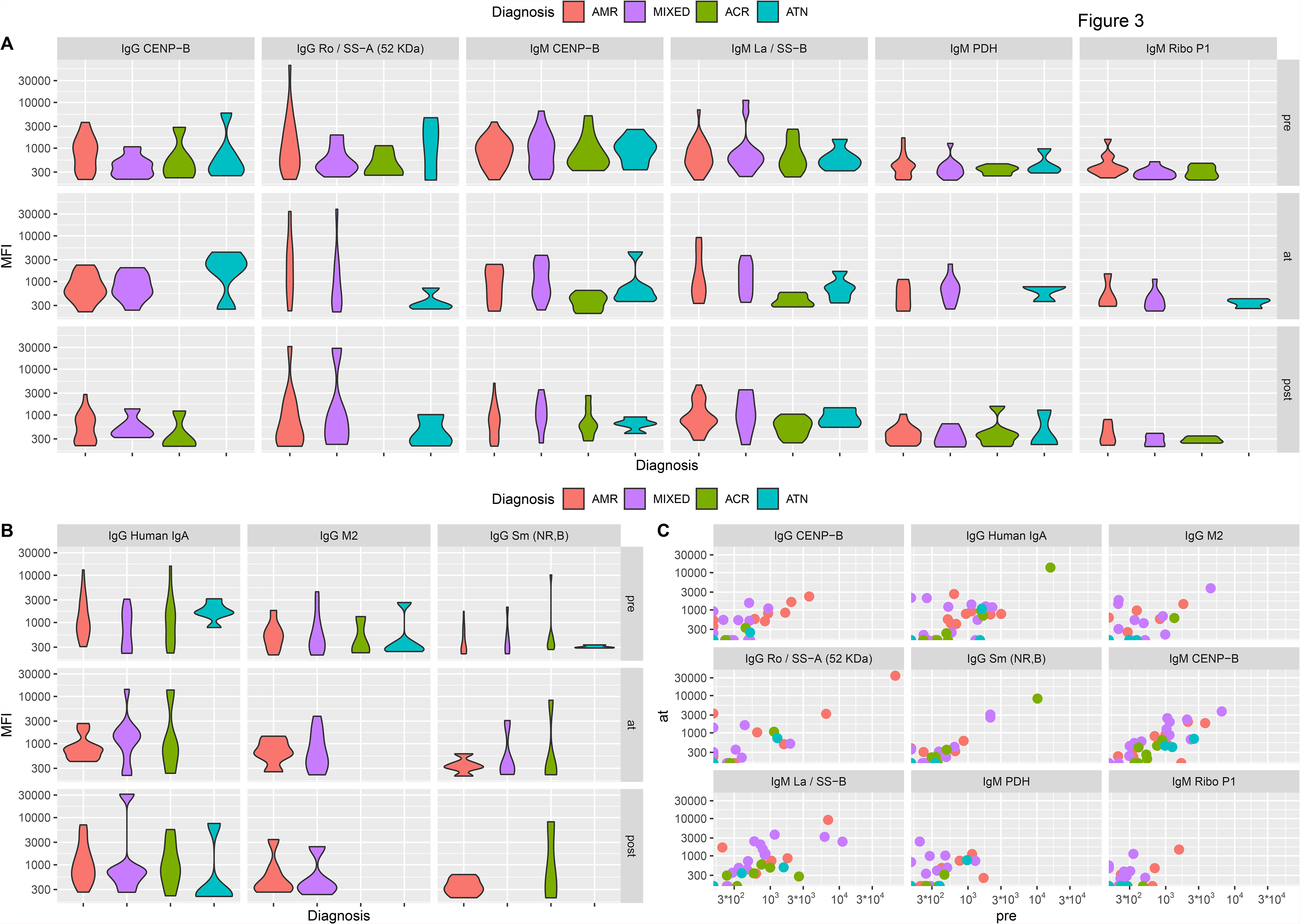
The evolution of top non-HLA antibodies increased in AMR and mixed rejection over time. The violin plots depict the distributions of the MFI values of the 6 antibodies significantly increased in AMR/mixed vs ACR (A) and the 3 antibodies significantly increased in AMR/Mixed vs ATN (B) at the time of diagnosis. Changes in the levels of these 9 antibodies over time were assessed by visualizing scatter plots of antibody MFI at diagnosis (y-axis) versus antibody MFI before transplant (x-axis), in patients who had both serum samples (C). AMR, antibody-mediated rejection; ACR, acute cellular rejection; ATN, acute tubular necrosis; Ig, immunoglobulin; MFI, median fluorescence intensity.

In addition to IgG anti-Ro/SS-A and anti-CENP-B, we found 17 non-HLA antibodies significantly altered before transplant. Fifteen of them were increased in AMR/mixed rejection (**Fig. S3**, **Table 3**). Pre-transplant levels of 7 IgG antibodies were significantly higher in AMR/mixed rejection than in ACR, including IgG against ribonucleoprotein SNRPD3, Histidine-tRNA ligase, and Desmin. Compared to ATN, AMR and mixed cases displayed significantly increased pre-transplant levels of 9 non-HLA antibodies, including anti-mitochondrial antibodies IgM anti-OGDC-E2 and IgG anti-PDH^61,62^. AMR/mixed patients also displayed higher pre-existing levels of IgM anti-LG3^49^, and increased antibodies against lysates of human glomerular microvascular endothelial cells (HGMEC) and cardiac endothelial cells (**Fig. S3**, **Table 3**). The distributions of these antibodies were similar between AMR and mixed cases (**Fig. S3**). Pre-transplant levels of 8 antibodies differed between sexes. While IgM against 3 ssDNA/dsDNA antigens were increased in women, men showed increased levels of IgG against Grp78/BiP, HGMEC lysate, and Asparaginyl-tRNA Synthetase (**Table S2**).

### Intraindividual Variability of Non-HLA Antibodies Over Time

We next evaluated antibody changes over time. We examined intraindividual changes in the levels of the 14 IgG and 11 IgM antibodies altered in AMR/mixed rejection pre-transplant and/or at diagnosis. For each antibody, we examined levels at diagnosis compared to pre-transplant. Most IgG antibodies did not fluctuate over time **(Fig. S4A**). This trend was also observed among IgM antibodies, although their intraindividual fluctuations were greater compared to IgG **(Fig. S4B**). Nonetheless, several AMR and mixed rejection patients displayed an increase in some antibodies at diagnosis, including IgG against Ro/SS-A(52KDa), human IgA, and M2, and IgM against La/SS-B, PDH, and Ribo P1 (**Fig. 3C**).

We next determined intergroup differences in antibody levels post-diagnosis. Twelve antibodies were significantly altered between groups. IgG against Troponin I, Thyroglobulin, dsDNA, and IgM anti-ADRB2 were significantly increased in AMR/mixed rejection, compared to ATN (**Table 3**, **Fig. S5**). Overall, non-HLA antibodies showed marked intraindividual consistency over time, but some antibodies increased specifically in AMR/mixed rejection at diagnosis.

### Non-HLA Antibodies Are Associated with Histopathology Features and DSA

We examined whether levels of non-HLA antibodies were associated with the presence of histopathological lesions and/or anti-HLA DSA. We focused on 26 non-HLA antibodies pre-transplant and 34 antibodies at diagnosis that were significantly and more strongly associated with at least one feature (P<0.05) (**Fig. 4**, **Table S3**).

**Figure 4.**
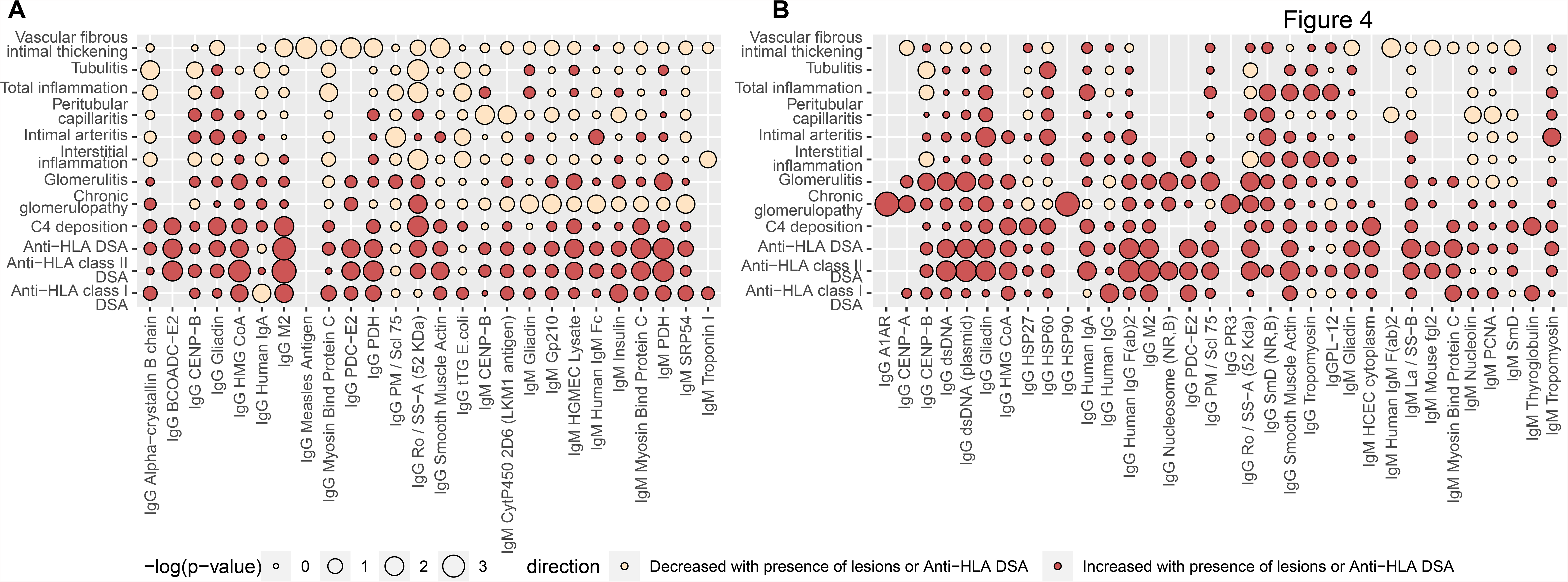
Association between levels of non-HLA antibodies and relevant histologic and clinical parameters. The bubble plot represents an association matrix between the presence of histopathology and serology features important in AMR and the MFI values of non-HLA antibodies before transplant (A) and at the time of diagnosis (B). The non-HLA antibodies that were more strongly associated with at least one clinical variable (according to p-value) are represented. The size of the nodes is inversely proportional to the p-value of the association. In turn, the color of the nodes indicates the direction of the association: increased antibody levels with the presence of histopathology lesions and/or presence of DSA are colored in red, while decreased antibody levels with the presence of histopathology lesions and/or presence of DSA are colored in beige. AMR, antibody-mediated rejection; ACR, acute cellular rejection; ATN, acute tubular necrosis; Ig, immunoglobulin; DSA, donor-specific antibodies; MFI, median fluorescence intensity.

Increased levels of IgG anti-Ro/SS-A(52KDa) and anti-CENP-B, both pre-transplant and at diagnosis, were significantly associated with the presence of peritubular capillaritis, glomerulitis, intimal arteritis, C4d deposition, and chronic glomerulopathy, but not with tubulitis or interstitial/total inflammation (**Fig. 4**). Increased pre-transplant levels of IgG and IgM anti-CENP-B were significantly associated with the presence of class-I and, more strongly, class-II DSA, while elevated IgG anti-Ro/SS-A(52KDa) was associated predominantly with the presence of class-II DSA (**Fig. 4A**). At diagnosis, increased levels of IgG anti-Ro/SS-A(52KDa) and IgG anti-CENP-B were significantly associated with the presence of class-I/II DSA (**Fig. 4B**). Accordingly, higher levels of antibodies pre-transplant and/or at diagnosis, but not post-diagnosis, tended to co-cluster with the presence of class-II DSA (**Fig. S2**). Two other antibodies increased in AMR/mixed rejection at diagnosis, namely IgM anti-La/SS-B and IgG anti-M2, were significantly associated with the presence of glomerulitis, C4d deposition, chronic glomerulopathy, and DSA (**Fig. 4B**). Interestingly, we found relevant clinical features were significantly associated with IgG against molecular chaperones: while higher IgG anti-HSP90 was linked to the presence of chronic glomerulopathy, increased IgG anti-HSP27 and anti-HSP60 levels were associated with C4d deposition and DSA (**Fig. 4B**). Few antibodies (6/26 pre-transplant and 13/34 at diagnosis), including IgG anti-dsDNA, IgG anti-HSP60, and IgG anti-Tropomyosin, were significantly and positively associated with tubulitis and/or total inflammation (**Fig. 4A**).

### External Validation of Antibodies against Ro/SS-A, CENP-B, and La/SS-B in AMR and Mixed Rejection

We next investigated if the observed increases in non-HLA antibodies in AMR/mixed rejection identified in the discovery group (Toronto) could be reproduced in an independent cohort. We conducted external validation of 6 antibodies significantly increased in AMR/mixed cases at diagnosis, namely IgG anti-Ro/SS-A(52 KDa), IgG and IgM anti-CENP-B, IgM anti-La/SS-B, IgM anti-PDH, and IgM anti-Ribo P1. We also interrogated their corresponding IgG or IgM levels (**Table S4**). We analysed serum samples from a previously described cohort of 60 kidney transplant patients (Montreal)^49^, including patients with AVR, ACR and stable recipients. We were able to reclassify a subgroup of AVR cases as having AMR or mixed rejection, or ACR grade 2-3. Concordantly with the discovery study, we observed significantly increased levels of IgG anti-Ro/SS-A(52 KDa, P=0.008) and IgG anti-PDH (P=0.02), and higher levels of IgG anti-La/SS-B (P=0.06), in AMR/mixed compared to ACR patients. When compared to stable controls, AMR/mixed patients still showed significantly increased levels of IgG anti-Ro/SS-A(52KDa, P=0.004) and IgG anti-La/SS-B (P=0.02), and higher levels of IgG anti-PDH (P=0.13) (**Fig. 5A-C**, **Table S4A**). The AMR/mixed group also displayed higher levels of IgM anti-CENP-B compared to ACR (P=0.06) and stable controls (P=0.05) (**Fig. 5D)**. Reassuringly, IgG anti-Ro/SS-A(52KDa) and IgM anti-CENP-B were significantly increased when comparing all cases with AVR with stable controls (P=0.02 and P=0.01, respectively), and remained elevated when compared to ACR (**Table S4B**).

**Figure 5.**
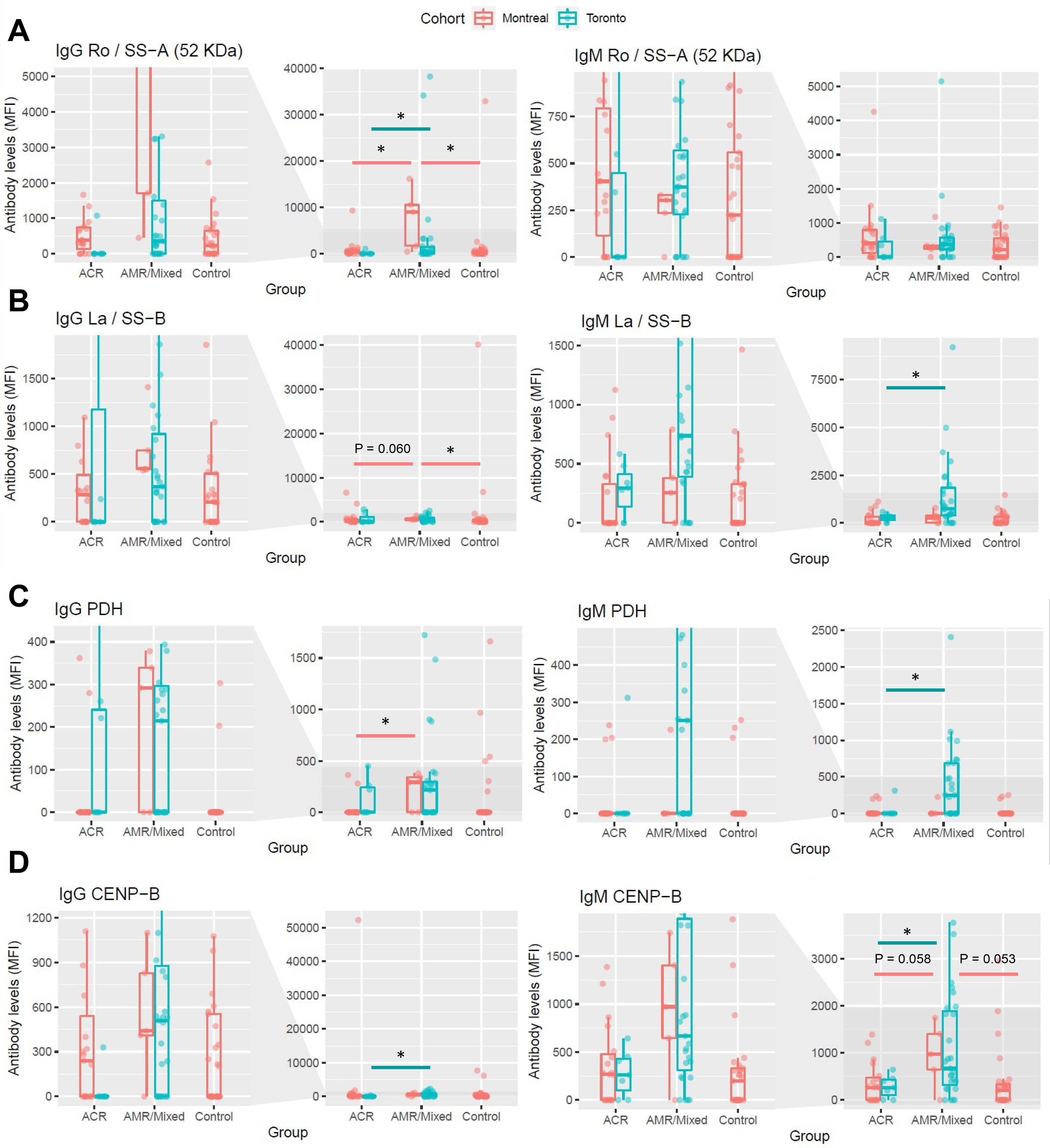
External validation of non-HLA antibodies increased in AMR and mixed rejection at the time of diagnosis. Differences between groups in the levels of four antibodies that were significantly increased in AMR/Mixed rejection at the time of diagnosis in the discovery cohort (Toronto), and were significantly associated with the presence of microvascular lesions, were interrogated in an external cohort (Montreal) for validation. For each antibody, levels from kidney transplant patients with AMR/Mixed rejection were compared to the levels from patients with ACR or stable non-rejecting kidney grafts (Control) and plotted next to their corresponding levels in the Toronto cohort. Levels of IgG and IgM against Ro/SS−A(52 KDa) (A), PDH (B), La/SS−B (C), and CENP-B (D) are shown. Data are represented as median ± interquartile range (IQR, box). *P<0.05 vs ACR or vs Control. AMR, antibody-mediated rejection; ACR, acute cellular rejection; MFI, median fluorescence intensity.

### Network Analysis Identifies Interactions Between Antibody Targets and Differentially Expressed Proteins in AMR

Antibodies against HLA and non-HLA antigens interact with proteins expressed by parenchymal cells, including endothelial and epithelial cells.^63–67^ We leveraged our recent proteomics study of glomeruli and tubulointerstitium in grafts with AMR compared to ACR and ATN,^48^ and built a protein-protein interaction network to study the connections between proteins significantly dysregulated in AMR kidneys, and protein targets of key antibodies identified in this study. We focused on targets of antibodies increased in AMR/mixed rejection and externally validated: TRIM21 (target of anti-Ro/SS-A(52KDa)), CENPB (target of anti-CENP-B), SSB (target of anti-La/SS-B), and PDHA1/PDHB (targets of anti-PDH) (**Fig. 6**). We found direct interactions between TRIM21, HSP90AB1 (increased in AMR glomeruli) and HLADRB1 (increased in AMR tubulointerstitium).^48^ The molecular chaperone HSP90AA1 and the proliferation marker PCNA connected with HLA class-I antigens (increased in AMR),^48^ and with antibody targets including centromeric proteins (CENPA, CENPB), metabolic enzymes (PDHA1, PDHB, DLAT, DBST, DBT), and ribosome-related proteins (SSB, RPLP1). The high connectivity between these proteins suggests biological relevance of both the proteins and antibodies directed against them in antibody-mediated injury.

**Figure 6.**
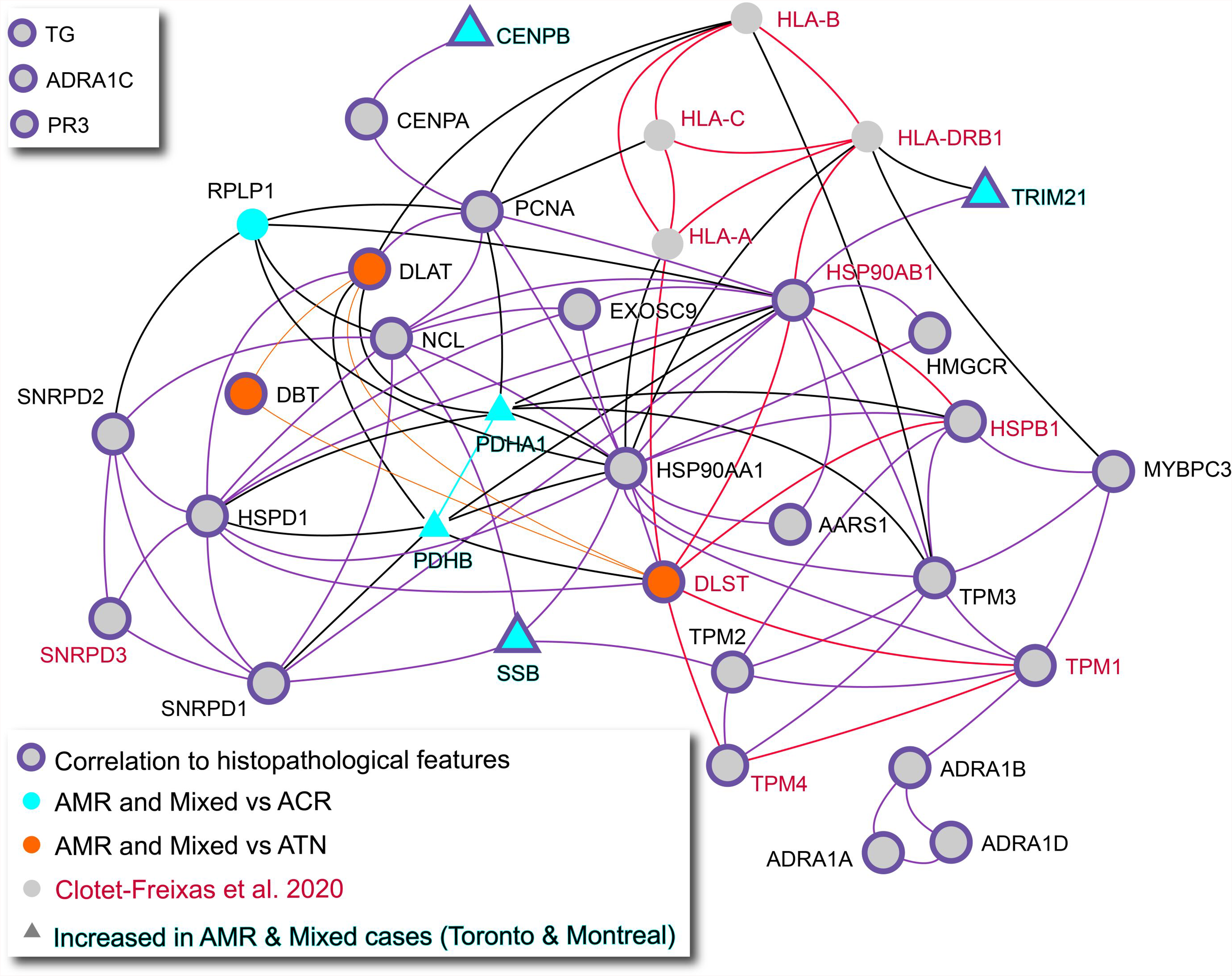
Network analysis of key antibody targets and proteins differentially expressed in kidney AMR. Physical protein-protein interactions of key non-HLA and HLA antibody targets were identified using the Integrated Interactions Database and visualized using NAViGaTOR 3.0.13. The selected targets were of relevance because their corresponding antibody was significantly increased in AMR/mixed patients at diagnosis, and/or significantly associated with the presence of AMR-related histopathology lesions or anti-HLA DSA. Turquoise and orange nodes represent the targets of non-HLA antibodies differentially increased in AMR/mixed patients compared to ACR and ATN, respectively. The nodes with purple highlight reflect targets of antibodies that were significantly associated with the presence of histopathology features and/or anti-HLA DSA. The gene names corresponding to the targets of antibodies validated in the external cohort (Montreal) are highlighted in turquoise. The nodes with a triangle shape represent targets of non-HLA antibodies increased in AMR/mixed rejection in both discovery and validation cohorts. The gene names of the non-HLA as well as HLA antibody targets that we previously found to be differentially expressed at the protein level in the AMR glomeruli or tubulointerstitium^48^ are colored in red. Purple edges connect proteins that are correlated to histopathological features, orange and turquoise edges connect proteins deregulated in AMR and mixed samples compared to ATN and ACR, respectively. Red edges connect proteins identified as deregulated in Clotet-Freixas *et al*., *JASN*, 2020. AMR, antibody-mediated rejection; ACR, acute cellular rejection; ATN, acute tubular necrosis.

## DISCUSSION

While autoantibodies against Ro/SS-A(52KDa), CENP-B and La/SS-B are elevated and pathogenic in several autoimmune diseases,^37,68–70^ their role in kidney allograft rejection has never been reported. Here, we show that 1) autoantibodies against Ro/SS-A(52KDa), CENP-B and La/SS-B were significantly higher in patients with AMR/mixed rejection compared to ACR at diagnosis; **2)** antibodies anti-Ro/SS-A(52KDa) and anti-CENP-B preceded transplantation and increased at the time of AMR/mixed diagnosis, in both early and late rejections; 3) these antibodies were associated with class-II DSA and microvascular lesions (**Fig. 7**). Our findings suggest that autoantibodies could participate in kidney allograft injury in AMR.

**Figure 7.**
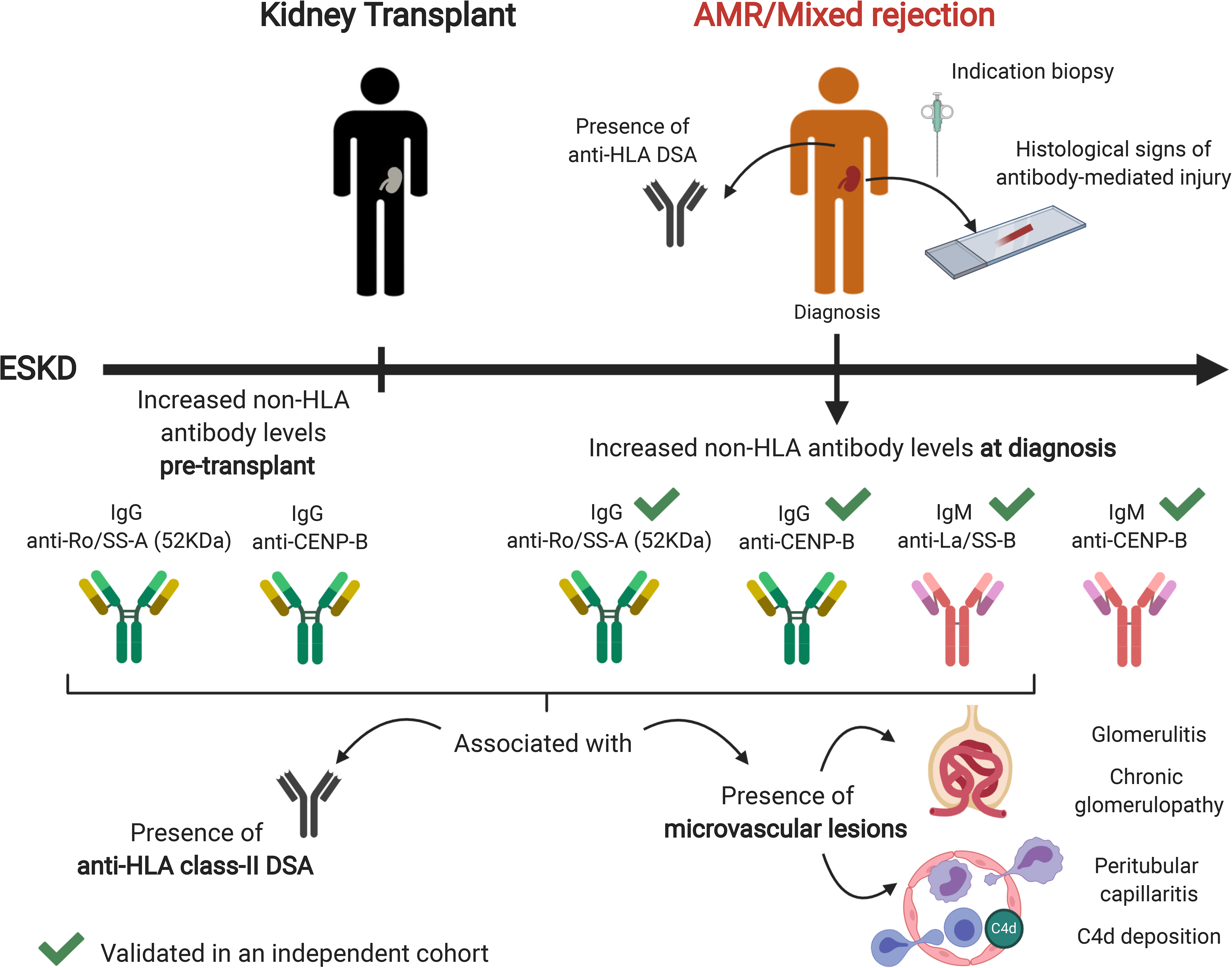
Summary of the key non-HLA antibodies associated with kidney AMR. Summary of relevant non-HLA antibodies increased in AMR/mixed rejection before transplant and at the time of diagnosis. IgG antibodies are depicted in green, while IgM antibodies are illustrated in red. The green ticks indicate that increased levels of IgG anti-Ro/SS-A(52KDa), IgG and IgM anti-CENP-B, and IgM anti-La/SS-B in AMR/Mixed rejection patients were validated in a second, independent cohort. ESKD, end-stage kidney disease; AMR, antibody-mediated rejection; Ig, immunoglobulin; DSA, donor-specific antibodies.

Despite similar distribution of total IgG and IgM levels between groups, our approach pinpointed specific antibodies significantly altered among different forms of allograft injury. In two independent cohorts, we demonstrated that autoantibodies against Ro/SS-A(52KDa), CENP-B and La/SS-B were increased in patients with AMR/mixed rejection compared to ACR, at diagnosis. Ro/SS-A(52KDa) antigen, also known as Ro52 or Tripartite motif-containing protein 21 (TRIM21), is recognized as the Sjögren’s syndrome-antigen A(SS-A)^71^, together with Ro60. Anti-Ro/SS-A antibodies have been described in autoimmune conditions including Sjögren’s syndrome, SLE, and systemic sclerosis, and proposed as markers of disease activity.^72^ TRIM21 is an Fc receptor that neutralizes opsonized viral particles entering cells^73^. TRIM21 can be upregulated and translocated to the nucleus under proinflammatory conditions, and modulate type-I interferon expression^74^. TRIM21 can also be expressed on the surface of apoptotic cells and become an immune target^75^. Monocyte surface TRIM21 expression was increased in patients with Sjögren’s syndrome, and upregulated by interferon-gamma^71^. Anti-TRIM21 antibodies specifically suppress the anti-inflammatory functions of this protein, while leaving type-I interferon production uncontrolled^76^. Anti-Ro antibodies could thus facilitate and enhance cytokine- and antibody-induced inflammation in AMR.

Anti-La/SS-B antibodies were elevated at diagnosis in AMR/mixed rejection. La/SS-B regulates cell cycle and binds to RNA polymerase-III transcripts, protecting them from exonucleases.^77^ Like TRIM21, La can be exposed on the surface of apoptotic cells, although it typically resides in the nucleus^75^. Positivity for both anti-Ro/SS-A and anti-La/SS-B antibodies has been observed in SLE and Sjögren’s syndrome.^37,68,78,79^ How these autoantibodies are generated is unknown. In mice, immunization with recombinant TRIM21 or La/SS-B resulted in loss of T-cell tolerance towards these antigens, and subsequent activation of B-cells to produce anti-Ro/SS-A and anti-La/SS-B antibodies^68^. The concomitant increase in anti-Ro/SS-A and anti-La/SS-B antibodies in our patients with AMR/mixed rejection is consistent with their similarities and associated phenotypes.

We also found significantly increased antibodies against major centromere autoantigen-B (CENP-B) in AMR/mixed rejection, compared to ACR. At diagnosis, both IgG and IgM anti-CENP-B were elevated in these patients. CENP-B is key to maintain chromosome segregation during mitosis^80^. CENP-B also binds to vascular cells stimulating proliferation, migration, and cytokine release.^81,82^ Anti-centromere antibodies have been described in several autoimmune and inflammatory diseases^39,70,83–85^. Senecal et al. demonstrated that anti-CENP-B antibodies inhibited proliferation and IL-8 production in vascular cells. Aberrant vascular repair and progressive arterial occlusion was observed in the presence of these antibodies^81^. We speculate that these antibodies may have similar effects in AMR.

Antibodies against Ro/SS-A(52KDa) and CENP-B preceded transplantation and increased at the time of AMR/mixed diagnosis, in both early and late rejections. Of note, only three patients in the AMR/Mixed rejection groups had SLE; thus, our observations are not related to pre-transplant autoimmune disease. While these antibodies were virtually absent in ACR, they were detectable in AMR/mixed cases, even before transplantation. Longitudinal sera enabled us to note that while all antibodies showed little variability between pre-transplant and at-diagnosis measurements, several antibodies including anti-Ro/SS-A(52KDa), anti-CENP-B and anti-La/SS-B increased at diagnosis compared to pre-transplant. This suggests that these antibodies, similar to anti-AT1R antibodies^11^, predated transplant and were formed by yet unrecognized mechanisms.

AMR and mixed rejection cases were diagnosed after distinct time intervals post-transplant. While pure AMR cases were biopsied within a month post-transplant, mixed rejection cases were diagnosed after a median of 174 days post-transplant. Despite this large difference in the time of diagnosis, the distribution of the anti-Ro/SS-A(52KDa), anti-CENP-B and anti-La/SS-B antibodies was remarkably similar in the two groups. This suggests that similar mechanisms are at play in AMR and mixed rejection, regarding the formation of these antibodies and their plausible influence on graft pathology.

Our third major observation was that increased levels of IgG anti-Ro/SS-A(52KDa), anti-CENP-B and IgM anti-La/SS-B were strongly associated with the presence of microvascular lesions and anti-HLA class-II DSA. Class-II DSA are more strongly associated with transplant glomerulopathy, and are considered to be more pathogenic than class-I^86–88^. However, we cannot rule out a stronger association with class-II DSA due to the higher prevalence of these antibodies compared to class-I in the AMR/mixed cohort. Along these lines, levels of IgG anti-Ro/SS-A(52KDa), anti-CENP-B and IgM anti-La/SS-B were associated with glomerulitis, C4d deposition and chronic glomerulopathy. Anti-CENP-B and anti-La/SS-B were also associated with intimal arteritis, linking the action of these antibodies in autoimmune diseases to lesions characteristic of AMR. As proposed for other non-HLA antibodies,^12,23^ these antibodies may act in synergy with anti-HLA DSA, enhancing allograft injury.

The targets of these non-HLA and HLA antibodies are highly interconnected in a protein-protein interaction network (**Fig. 6**). Furthermore, integration with our recent kidney tissue proteomics dataset^48^ highlighted that proteins perturbed in the AMR tissue are directly connected with the targets of the immune response. The key hubs in this network are chaperones HSP90, which participate in renal immunity^89^. HSP90 was previously elevated in the serum of patients with kidney AMR^90^. We demonstrated that anti-HSP90 antibodies were strongly associated with the presence of chronic glomerulopathy. PCNA is another hub in the network, connecting CENP, PDH and HSP90 proteins. Upon injury, increased PCNA expression indicates increased cell cycle entry, which may lead to adverse events such as hypertrophy or mitotic catastrophe.^91,92^ In turn, antigen presentation and HLA-ligation can trigger proliferation in endothelial cells.^87,93^ Proliferative stress in kidney AMR may result in abnormal centromere function and affect the turnover of centromere-related proteins, such as CENP-B. In conclusion, proteins disrupted in kidney tissue during AMR interact directly with the targets of anti-HLA and non-HLA antibodies. Further studies aimed at deciphering the biology of these antibodies and their target proteins is warranted.

Our study has several strengths. We investigated a sizeable group of well-characterized kidney transplant recipients, with longitudinally-collected serum samples. Using an innovative protein microarray, we studied non-HLA antibody changes in different subgroups and over time. We integrated the findings from this study with clinical/histopathologic data and with our prior proteomics-based study.^48^ The key findings were externally validated. Our study also has limitations. After excluding sera affected by PLEX and/or IVIG, the study of antibody dynamics was limited to a smaller subset of patients. In addition, the time intervals between the indication biopsy date and the post-diagnosis serum were extremely variable, preventing us from including this time point in the study of dynamic changes in antibody levels. Finally, the current study pinpoints novel and interesting associations, but further work is required to establish their causal relationship with AMR.

In conclusion, our approach revealed a novel link between increased pre-transplant levels of IgG anti-Ro/SS-A(52KDa) and anti-CENP-B and the development of AMR after kidney transplantation. Together with IgM anti-La/SS-B, these antibodies were increased in AMR early and late after transplant. All 3 antibodies were associated with the presence of microvascular lesions and anti-HLA class-II DSA, suggesting that they may synergize with class-II DSA and induce endothelial injury in AMR. This is the first effort to date that links specific pre-transplant non-HLA autoantibodies with the diagnosis of AMR after kidney transplantation.

## DISCLOSURES

All authors have nothing to disclose.

Dr. Igor Jurisica reports receiving personal fees from Canadian Rheumatology Association, grants and nonfinancial support from IBM, and personal fees from Novartis, outside the submitted work.

## FUNDING

AK is supported by Kidney Foundation of Canada Predictive Biomarker grant KFOC160010, the Canadian Institutes of Health Research (CIHR), Canada Foundation for Innovation (CFI) grant 37205, and Kidney Research Scientist Core Education and National Training (KRESCENT) program grants CIHR148204, KRES160004, and KRES160005. AK has also received funding from the Toronto General and Western Hospital Research Foundation (TGTWF 1617-464; TGTWF MKFTR 1718-1268). SC-F is supported by the KRESCENT program (2019KPPDF637713). IJ, CP, and MK were supported in part by Ontario Research Fund grant 34876, Natural Sciences and Engineering Research Council of Canada grant 203475, and CFI grants 29272, 225404, and 30865. HC is a Fonds de recherche du Québec (Santé) Junior 2 scholar.

## ACKNOWLEDGEMENTS

Special thanks to Marc Angeli, Sharon Selvanayagam, and Dr. Uzma Nadeem.

## AUTHOR CONTRIBUTIONS

Dr. Ana Konvalinka conceived the study.

Dr. Ana Konvalinka, Dr. Rohan John, Dr. Andrzej Chruscinski, Dr. Sergi Clotet-Freixas, Dr. Heloise Cardinal, and Dr. Mélanie Dieudé participated in study design.

Dr. Ana Konvalinka, Dr. Caitriona McEvoy, Dr. Sonia Rodríguez-Ramírez, and Dr. Sergi Clotet-Freixas retrieved and curated clinical data from the discovery cohort.

Dr. Heloise Cardinal, Dr. Mélanie Dieudé, and Dr. Marie-Josée Hébert retrieved and curated clinical data from the validation cohort.

Dr. Andrzej Chruscinski and Dr. Sergi Clotet-Freixas performed the experiments.

Dr. Chiara Pastrello, Dr. Max Kotlyar, Dr. Igor Jurisica, Dr. Ana Konvalinka, Dr. Caitriona McEvoy, Dr. Sergi Clotet-Freixas, and Dr. Sofia Farkona analyzed the data.

Dr. Chiara Pastrello, Dr. Max Kotlyar, Dr. Sonia-Rodríguez-Ramírez, and Dr. Sergi Clotet-Freixas made the figures.

Dr. Sergi Clotet-Freixas, Dr. Sonia-Rodríguez-Ramírez, and Dr. Ana Konvalinka drafted and revised the paper; and all authors approved the final version of the manuscript.

Dr. Caitriona M. McEvoy, Dr. Yanhong Li, Peixuen Chen, and Emilie Chan retrieved and curated clinical data; Dr. Yanhong Li, Dr. Olusegun Famure, and Dr. S. Joseph Kim selected the cases from the CoReTRIS registry and performed case-control matching.

## SUPPLEMENTAL MATERIAL - TABLE OF CONTENTS

**Supplemental Table 1.** List of non-HLA antigens studied in the protein array.

**Supplemental Table 2.** Pre-transplant antibodies against non-HLA antigens significantly altered between sexes.

**Supplemental Table 3.** Study of the association between IgG and IgM non-HLA antibody levels and relevant clinical and histological variables.

**Supplemental Table 4.** External validation of key non-HLA antibodies.

**Supplemental Figure 1.** Distribution of antibody levels according to patient sex.

**Supplemental Figure 2.** Clustering analysis of non-HLA antibodies significantly altered in AMR and mixed rejection.

**Supplemental Figure 3.** The evolution over time of 17 non-HLA antibodies increased in AMR and mixed rejection before transplant.

**Supplemental Figure 4.** The changes in the levels of relevant non-HLA antibodies over time, among patients who had both pre-transplant and at-diagnosis sera.

**Supplemental Figure 5.** The evolution over time of 11 non-HLA antibodies increased in AMR and mixed rejection after diagnosis.

---

**Figure S1.**
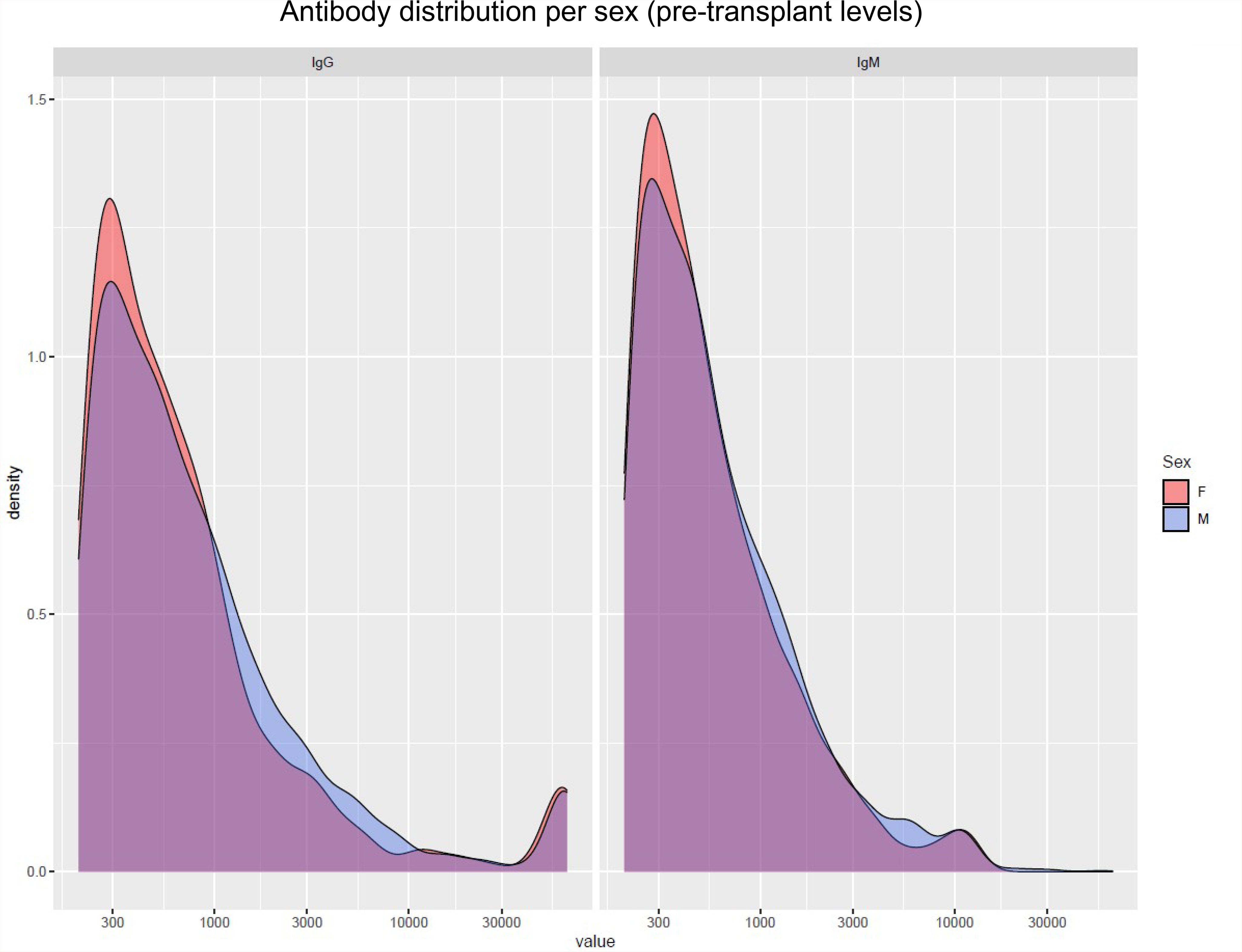
Distribution of antibody levels according to patient sex. The density plots depict the distributions of IgG and IgM MFI values before transplant in male (M, purple) and female (F, orange) patients. In each patient, a value of zero was given to all antibodies with MFI < 200. Log2-transformed MFI values were used to create the plots using ggplot2 3.3.2 in R. The x-axis encompasses the range of all MFI values among the detected antibodies, while the y-axis represents the frequency (density) of each of these values. AMR, antibody-mediated rejection; ACR, acute cellular rejection; ATN, acute tubular necrosis; Ig, immunoglobulin; MFI, median fluorescence intensity.

**Figure S2.**
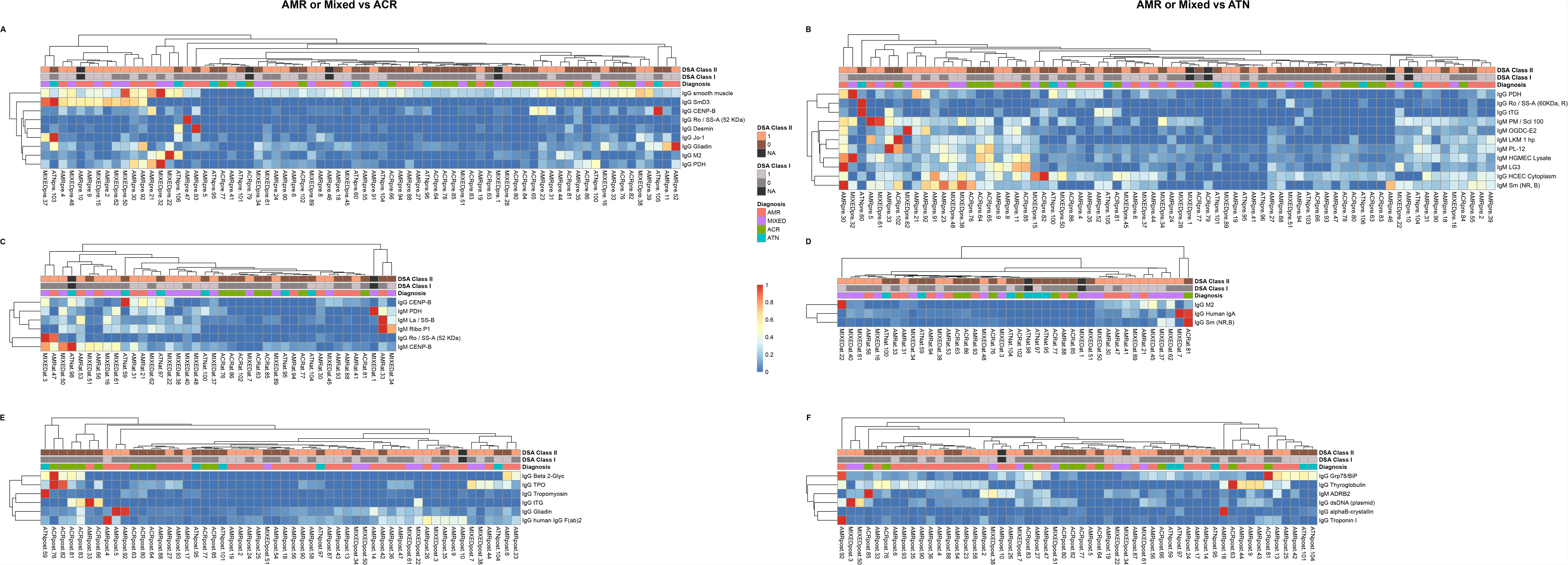
Clustering analysis of non-HLA antibodies significantly altered in AMR and mixed rejection. The heatmaps illustrate the hierarchical clustering analyses of the non-HLA antibodies significantly differentially increased or decreased in AMR and mixed rejection before transplant (A,B), at the time of diagnosis (C,D), and post-transplant (E,F), compared to ACR or ATN (P<0.05). Log2-transformed MFI values of each antibody were used to build the heatmaps. For each antibody, a value of zero was given to the samples with MFI < 200 (below the limit of detection). Patient clustering was evaluated according to their diagnosis and the presence/absence of anti-HLA class-I and/or anti-HLA class-II DSA. AMR, antibody-mediated rejection; ACR, acute cellular rejection; ATN, acute tubular necrosis; Ig, immunoglobulin; DSA, donor-specific antibodies; MFI, median fluorescence intensity.

**Figure S3.**
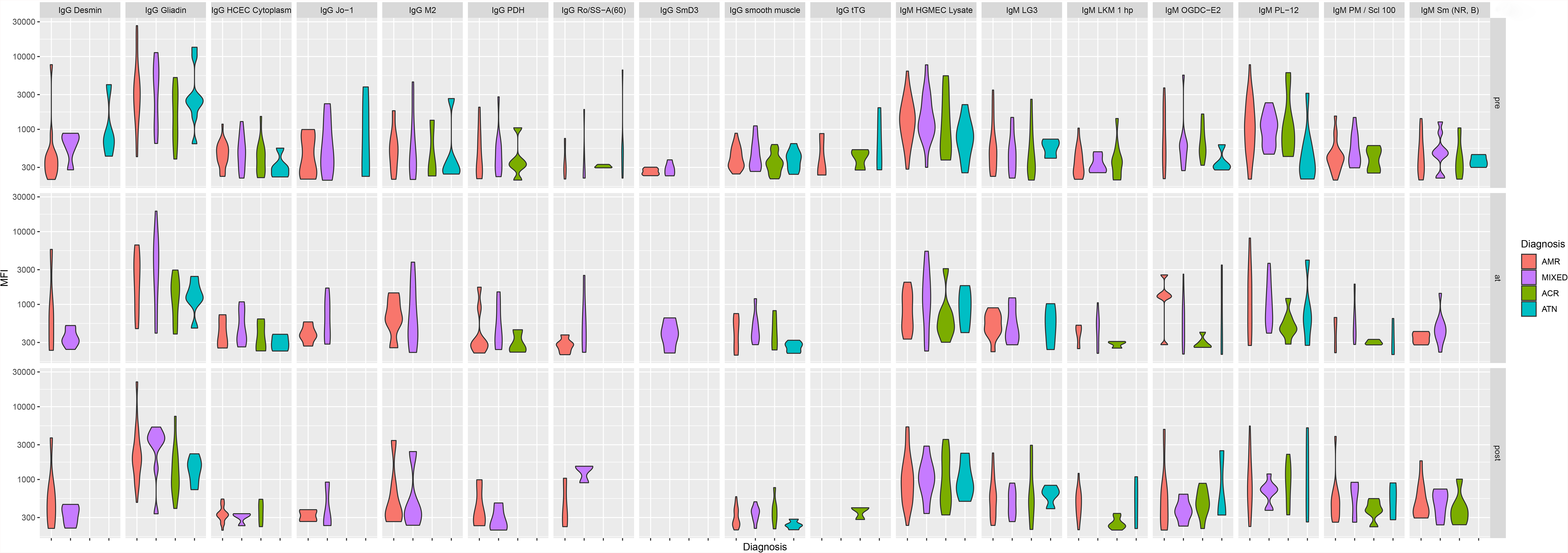
The evolution over time of 17 non-HLA antibodies increased in AMR and mixed rejection before transplant. The violin plots depict the distributions of the MFI values of 17 antibodies significantly altered before transplant in AMR/Mixed patients compared to ACR and/or ATN. AMR, antibody-mediated rejection; ACR, acute cellular rejection; ATN, acute tubular necrosis; Ig, immunoglobulin; MFI, median fluorescence intensity.

**Figure S4.**
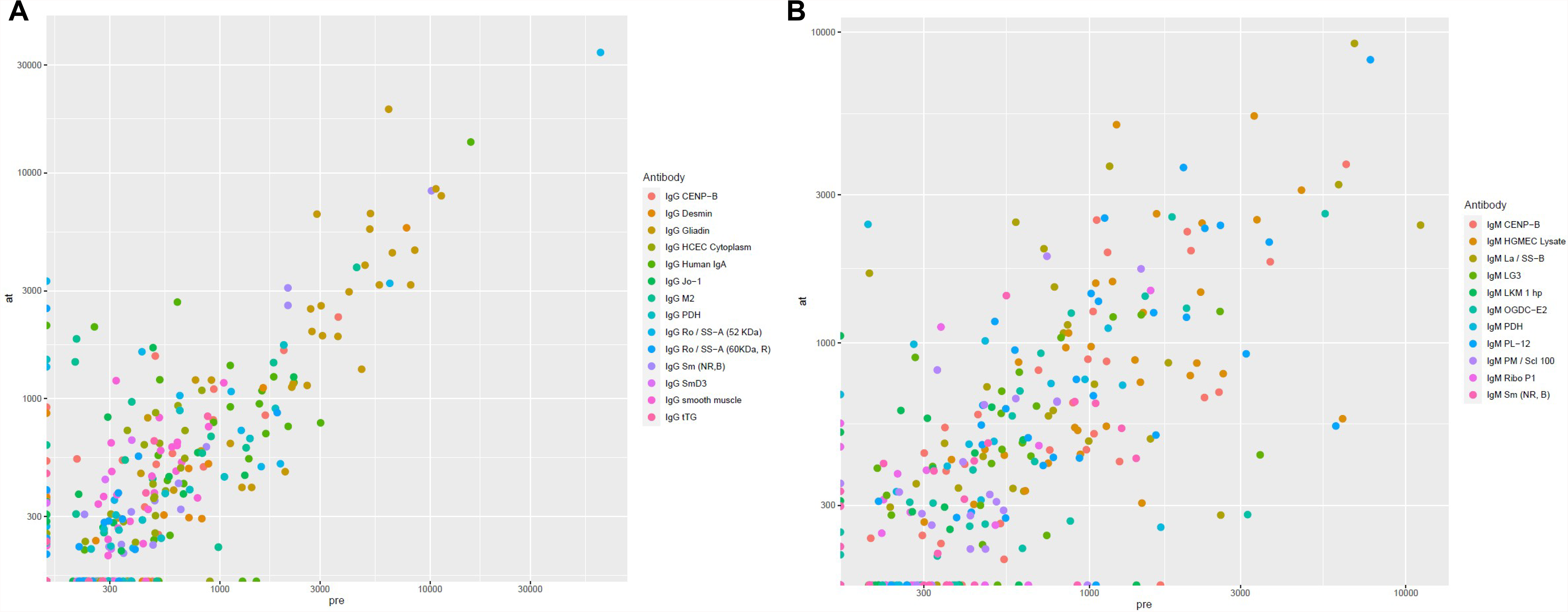
The changes in the levels of relevant non-HLA antibodies over time, among patients who had both pre-transplant and at-diagnosis sera. Changes over time in the levels of the 14 IgG antibodies (A) and 11 IgM antibodies (B) altered in AMR/mixed rejection before transplant and/or at the time of diagnosis were studied. Antibody fluctuations were assessed by visualizing antibody MFI at diagnosis (y-axis) versus antibody MFI before transplant (x-axis) using scatter dots. Overall, most antibodies did not appear to fluctuate in individuals over time. AMR, antibody-mediated rejection; Ig, immunoglobulin; MFI, median fluorescence intensity.

**Figure S5.**
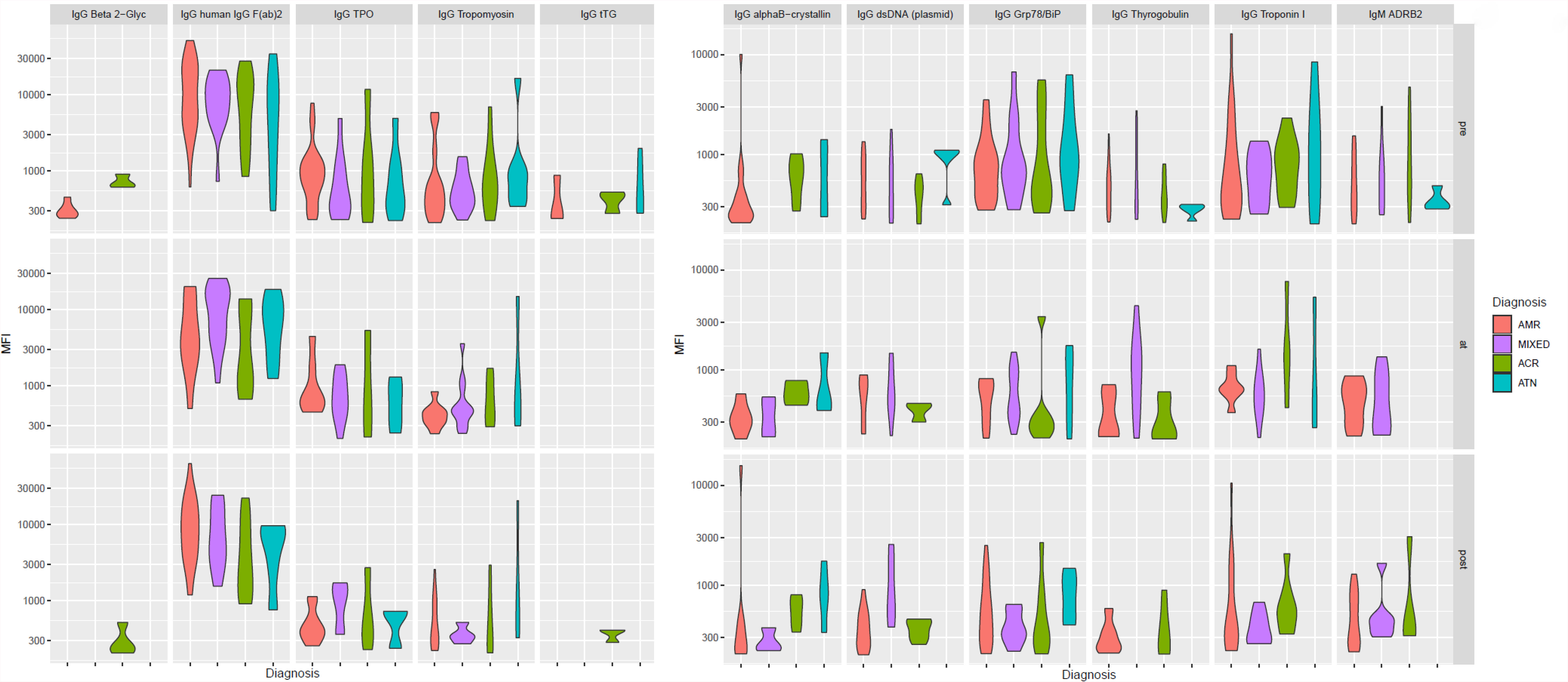
The evolution over time of 11 non-HLA antibodies increased in AMR and mixed rejection after diagnosis. The violin plots depict the distributions of the MFI values of 11 antibodies significantly altered after diagnosis in AMR/mixed patients compared to ACR and/or ATN. AMR, antibody-mediated rejection; ACR, acute cellular rejection; ATN, acute tubular necrosis; Ig, immunoglobulin; MFI, median fluorescence intensity.

